# Microstructural heterogeneity of white-matter aging and its age-dependent coupling to cognition

**DOI:** 10.64898/2026.10.09.757817

**Authors:** Rowena Chin, Xiaoyang Hu, Jasmine Ranieri, Ji-Hyun Kim, Hwamee Oh

## Abstract

Brain white matter (WM) microstructure undergoes extensive changes with age, yet the regional and cellular heterogeneity of age-related WM alterations and their impact on cognition remain unclear. Leveraging Neurite Orientation Dispersion and Density Imaging (NODDI), an advanced microstructural modelling to capture specific WM cellular properties, we investigated age-dependent relationships between advanced microstructural markers and cognitive performance, indexed by six cognitive domains derived from principal component analysis. Using a suite of standard diffusion tensor imaging (DTI) and NODDI techniques in eighty-six cognitively normal adults (Young: n=33, Older: n=53), we show that age group differences are widespread across whole brain white matter, with converging region-based and voxel-level evidence implicating commissural, association, and limbic pathways. Older adults exhibited significant and widespread reductions in global fractional anisotropy (FA) and neurite density index (NDI), concomitant with increases in mean diffusivity (MD), orientation density index (ODI), and free water fraction (FWF). Critically, age moderated WM-cognition associations in a domain- and tract-specific manner: younger adults exhibited stronger coupling between commissural and association tract microstructure and language and visual memory, whereas older adults showed greater dependence on projection and limbic tracts for executive and language function. The observed age-dependent coupling between specific microstructural metrics and cognition underscores the value of advanced diffusion models for capturing heterogeneity in brain aging, reveals age-specific biomarkers of cognitive health. These patterns align with phylogenetic and ontogenetic patterns of white matter vulnerability in callosal, long-range association, and limbic pathways, and suggest cellular specificity in white matter changes associated with cognitive decline.

**Significance Statement:** Aging affects brain white matter in complex and regionally specific ways, yet the biological bases of these changes and their relevance to cognition remain challenging to disentangle. Using multi-parameter diffusion MRI modeling, we found that age-related decline follows organized spatial patterns, with the greatest vulnerability in long-range association, callosal, and limbic pathways. These regional effects arise from distinct microstructural processes, including widespread extracellular alterations and more localized changes in neurite density and fiber organization. The strength and direction of white matter–cognition relationships differ between younger and older adults, revealing age-dependent shifts in how brain structural connections support behavior. Together, these findings identify biologically meaningful dimensions of white-matter vulnerability that may explain individual differences in cognitive aging.

## Introduction

Brain white matter (WM) forms the connective architecture for efficient neural communication and underpins a broad range of cognitive functions (Fields, 2008). With advancing age, WM microstructure undergoes progressive reorganization and loss of integrity, contributing to age-related cognitive decline (Salat et al., 2005; Madden et al., 2012). Aging of cerebral WM reflects multiple concomitant neurobiological processes, including demyelination (Marner et al., 2003), axonal degeneration (Salvadores et al., 2017), neuroinflammation (Raj et al., 2017), and tissue loss, even after accounting for the presence of cortical atrophy (Guttmann et al., 1998). These cellular factors manifest macroscopically as reductions in directional organization and increased water mobility within tissue. Diffusion-weighted magnetic resonance imaging (dMRI) has been widely used to characterize these microstructural changes *in vivo* and has consistently demonstrated reduced fractional anisotropy (FA) and increased mean diffusivity (MD) (Bennett et al., 2010). These effects often follow reproducible spatial gradients of decline, most prominently along anterior-posterior and superior-inferior axes (Head et al., 2004), suggesting that WM aging is not spatially uniform but follows organized topographical patterns linked to tract type vulnerability and myelination timing (Bartzokis, 2004). Within these gradients, long-range association pathways, prefrontal callosal fibers, and certain projection and limbic pathways such as the thalamic radiations and fornix, have shown disproportionate susceptibility in aging.

Beyond mapping structural decline, diffusion imaging has been instrumental in linking WM microstructure integrity to cognitive function in vivo. Across healthy and clinical populations, DTI studies extensively demonstrate that reduced anisotropy and elevated diffusivity are associated with poorer cognitive performance (Bennett et al., 2010; Madden et al., 2012). Both global and regional effects have been described. Global whole-brain measures of white matter integrity have been linked to broad cognitive abilities such as processing speed and executive function (Madden et al., 2009; Vernooij et al., 2009), whereas more focal associations have been observed for specific cognitive domains, for example, memory performance with limbic and temporal lobe tracts including the fornix and cingulum (Charlton et al., 2010; Metzler-Baddeley et al., 2011; Archer et al., 2020). Notably, many of these frontotemporal association, callosal, limbic pathways that relate to cognitive performance in healthy adults also show early diffusion abnormalities in individuals with mild cognitive impairment or at elevated risk for Alzheimer’s disease (Sexton et al., 2011), a convergence that may represent a continuum between normative aging and prodromal neurodegeneration.

Despite the significant association between global and regional white matter aging and age-related cognitive decline, it remains challenging to disentangle biological underpinnings of the contribution of white matter aging to age-related cognitive changes with greater specificity. Multi-compartment diffusion models such as Neurite Orientation Dispersion and Density Imaging (NODDI) address this limitation by decomposing the diffusion signal into tissue compartments – neurite density (NDI), indexing intra-axonal packing and myelin content; orientation dispersion (ODI), reflecting variability of fiber orientation and coherence, and free-water fraction (FWF), representing extracellular diffusion arising from cerebrospinal and interstitial fluid, which collectively offers greater biological specificity than conventional tensor metrics (Zhang et al., 2012). Aging studies generally report decreased NDI and increased ODI and FWF in older adults, consistent with diminished axonal density, greater fiber disorganization, and expansion of the extracellular compartment due to vascular, inflammatory, or atrophic processes (Pasternak et al., 2012; Merluzzi et al., 2016). Although prior work has established that WM microstructure declines with age and is related to memory and executive function test performance, less is understood about how different putative cellular components of white matter account for age-related cognitive changes across a wide range of cognitive domains. It also remains unknown whether aging alters how WM supports cognition, i.e. whether the same biological features of WM support cognition to the same extent in younger and older adults, and if these potential age-dependent structure-behavior couplings are generalizable across spatial scales, from focal tracts to global WM architecture.

The present study addresses these questions using a multi-parameter diffusion MRI approach in a cohort of healthy younger and older adults from the Multimodal Imaging of Cognition, Aging, and Alzheimer’s Disease (MICAAD) Study. We combined tensor and NODDI modeling to characterize age effects on WM microstructure across global skeleton, regional tract-based, and voxel-wise scales, and to test whether microstructural measures show age-dependent associations with cognition across data-driven cognitive domains. By jointly leveraging conventional DTI indices and advanced NODDI parameters, we aimed to delineate the microstructural components underlying healthy WM aging and to capture age-linked alterations in structure-behavior relationships. We hypothesized that older adults would show widespread reductions in FA and NDI and increases in MD, ODI, and FWF, reflecting both intracellular and extracellular components of tissue degeneration, and that age would moderate WM-cognition relationships. This work provides a mechanistic, *in vivo* account of how aging alters both the microstructural organization of WM and clarifies its role in shaping age-related differences in cognitive performance.

## Methods

### Study cohort

Eighty-six participants (Young: n=33; Older: n=53) who performed a multi-shell diffusion weighted MRI scan from the Multimodal Imaging of Cognition, Aging, and Alzheimer’s Disease (MICAAD) Study was included in the present study. The MICAAD study, which was launched in 2020, is a longitudinal observational study designed to investigate the cognitive, affective, and neural changes in association with brain aging and Alzheimer’s disease pathologies. A total of one hundred eighty-three participants consisting of 61 healthy young (mean age=25.1 years old (SD=6.9), 38 females) and 122 clinically intact older adults (mean age=68.5 years old (SD=5.7), 80 females) were enrolled in the study at baseline. Healthy young (18–35 years) and clinically intact older adults (≥60 years) were recruited from the community via on-line advertisements, community newsletters or emails, and flyers. Participants were initially screened by telephone, following which they came to the laboratory and underwent a medical interview and a detailed battery of neuropsychological tests, including the National Alzheimer’s Coordinating Center’s Uniform Data Set (UDS3) battery. All participants were required to be fluent in English, have MMSE scores ≥ 26, Clinical Dementia Rating scale (CDR) = 0, and have at least a fourth grade reading level. Participants were screened for MRI contraindications and hearing or visual impairment that would impede testing. In addition, participants must meet the following criteria: no objective cognitive impairment; no history of significant neurological disease (including stroke, seizure, loss of consciousness); no current diagnosis of severe psychiatric disorder, no significant systemic medical illness; no use of medications affecting cognition; no MRI or PET contraindications; and not currently pregnant. Participants were also excluded if they had uncontrolled high blood pressure (systolic blood pressure ≥180mmHg or diastolic blood pressure ≤ 105mmHg). All participants provided written informed consent in accordance with the Declaration of Helsinki prior to their participation and the study protocol was approved by the Institutional Review Board at Brown University. All participants were compensated for participation. The current analyses were drawn from the subset of the MICAAD participants who completed baseline diffusion MRI and neuropsychological assessments.

### Neuropsychological assessment

We performed principal component analysis (PCA) on a subset of the MICAAD study (138 participants) to develop a more concise and reliable summary measure from multiple neuropsychological test measures. This included 82 cognitively intact older adults (mean age, 68.7 years old (SD=5.9); range, 60–90 years; 59 females; mean Mini Mental State Examination [MMSE], 29 (SD=1.2)) and 56 young adults (mean age, 24.4 years old (SD=5.2); range, 18–35 years; 37 females; mean MMSE, 29.1 (SD=1.2)). All subjects included in PCA completed 22 cognitive performance test measures. The tests included Immediate and Delayed Recall of the UDS Craft Story 21, Benson Complex Figure Immediate and Delayed Copy (Delis et al., 2016), Stroop test (Golden, 1978), “Trail A, Trail B, and Trail B minus A” from Trail Making Test (TMT) A and B (Reitan, 1958), Digit Symbol subtest (Wechsler, 1997), category “Vegetables” and “Animals” from Category Fluency Test (Benton et al., 1983). Digit span forward and backward, total scores of the Multilingual Naming Test, Verbal fluency of letters F and L, total recall and delayed recall of the Selective Reminding Test (SRT; Buschke and Fuld, 1974), Letter Number sequencing, and the American National Adult Reading Test (AMNART; Grober and Sliwinski, 1991) as an estimate of IQ.

PCA is a multivariate method that extracts underlying, unobserved components among observed variables that are correlated with each other to varying degrees, resulting in a reduction of a large number of variables into a smaller number of variables (Tabachnick and Fidell, 2001). First, cognitive test scores were entered into a PCA with varimax rotation. In order to ensure that cognitive test scores included in PCA are correlated with each other enough for components to emerge, the correlation matrix of cognitive test scores was examined to verify that a majority of correlations exceed R= 0.30 (Tabachnick and Fidell, 2001). Components were extracted with a criterion of an eigenvalue greater than 1 (Kaiser, 1960). Based on the individual tests that expressed the highest component loading scores for the component, each component was named as follows: executive function (EXE), language (LAN), verbal episodic memory (VerMEM), semantic memory (SM), working memory (WM), and visual episodic memory (VisMEM). Each subject’s score for each component was determined by regression of a subject’s raw test scores into the component loading scores for the component. PCA was performed in R (v4.3.1).

### Imaging data acquisition

Neuroimaging data were acquired on a 3T Siemens PRISMA MRI system equipped with a 64-channel head coil at the Imaging Research Facility at Brown University. Diffusion-weighted imaging employed a multi-shell, spin-echo echo-planar imaging (EPI) sequence with the following parameters: repetition time (TR) = 3400 ms; echo time (TE) = 71 ms; flip angle = 180°; field of view (FOV) = 232 mm; resolution = 2 mm isotropic, acquisition time = 7 min 25 sec. The acquisition included twelve b=0 s/mm² images and diffusion-weighted volumes at b = 500, 1000, and 2000 s/mm², with 114 gradient directions. Additional reversed phase-encoded b=0 images were acquired to support post-acquisition susceptibility distortion correction.

### Imaging data processing

#### Diffusion MRI preprocessing

Diffusion MRI preprocessing was performed using FSL v6.0.5 (FMRIB Software Library, Oxford, UK). Susceptibility distortions were corrected using FSL’s topup, applied to opposing phase-encoded b=0 volumes. Brain extraction was conducted with threshold of 0.2 and brain masks were visually inspected. Motion and eddy current distortions were corrected using FSL’s eddy, which included slice-to-volume motion correction, outlier replacement, and field map integration. Following preprocessing, voxel-wise diffusion tensor estimation was conducted using FSL’s dtifit, which generated whole brain white matter microstructural maps of fractional anisotropy (FA) and mean diffusivity (MD) in native space.

#### NODDI microstructure modeling

To derive a more comprehensive range of white matter microstructural markers, we applied the Accelerated Microstructure Imaging via Convex Optimization (AMICO) implementation of the Neurite Orientation Dispersion and Density Imaging (NODDI) model (Daducci et al., 2015). AMICO was selected for its computational efficiency and stability in large-scale datasets. Preprocessed diffusion data were converted into the required format, including gradient directions, b-values, and acquisition scheme files. The default three-compartment model was used to estimate neurite density index (NDI), orientation dispersion index (ODI), and free-water fraction (FWF). Outputs were retained in native space prior to normalization.

#### Spatial normalization

All diffusion-derived maps (FA, MD, NDI, ODI, FWF) were spatially normalized using the Tract-Based Spatial Statistics (TBSS) pipeline (Smith et al., 2006). Each subject’s FA image was nonlinearly registered to the FMRIB58_FA template. These deformation fields were then applied to the corresponding MD, NDI, ODI, and FWF maps. All normalized images were concatenated into group-level volumes and manually reviewed for quality control.

#### Global mean white matter measures

Global white matter metrics were computed by projecting each subject’s diffusion-derived maps onto the TBSS mean white matter skeleton and a threshold of 0.3 was applied to retain the central portions of major white matter tracts. For each diffusion modality (FA, MD, NDI, ODI, FWF), the mean value within the thresholded skeleton was extracted for each subject, yielding a summary index of whole brain white matter microstructure per modality.

#### Regional tract based white matter measures

Region-specific analyses were conducted using the JHU ICBM-DTI-81 white matter labels atlas which defines 50 whole brain, non-overlapping white matter tracts in MNI space (Wakana et al., 2004). Individual binary masks were generated for each tract region-of-interest (ROI), and mean white matter microstructural measures (FA, MD, NDI, ODI, and FWF) were extracted across all ROIs for each subject.

### Statistical analysis for global and tract-based averaged white matter microstructure measures

All statistical analyses were performed in R (v4.2.3). Permutation-based general linear modeling was implemented using the *permuco* package (v1.1.0) with the Freedman–Lane method and 5000 permutations to obtain non-parametric p-values for each model term.

We first examined age group effects on white matter microstructure, including sex as a covariate, leveraging data from the full sample cohort of 86 participants. Linear regressions were conducted separately for each combination of microstructural measure (FA, MD, NDI, ODI, FWF) at two analysis levels: (i) a global summary measure derived from the thresholded mean white matter skeleton, and (ii) regional values from each atlas-defined tract of interest. In addition to the age group effect on microstructural measures, associations between chronological age (years) and global summary mean white matter measures were evaluated using Spearman’s rank-order correlations, given non-normality of at least one variable in all cases as determined by the Shapiro–Wilk test.

The second set of regression models investigated the effect of white matter microstructure on cognitive performance across different cognitive domains in a subset of participants with complete neuropsychological data (n=73). Age group was included as a categorical variable, as well as an interaction term of age group and microstructure measures, with sex and years of education as covariates. Cognitive performance was represented as six PC scores, each corresponding to a distinct cognitive domain. As in the primary analysis, models were performed in two-level runs. Specifically, this meant distinct models were estimated for the effect of each microstructural measure using either global whole brain average or regional tract-specific metric and its interaction effect with age group on cognitive domain factor score.

To quantify the magnitude of statistically significant effects, partial eta-squared (η²p) was calculated from tract ROI-level permutation-based GLM analyses. As permutation tests do not directly provide effect sizes, the F-statistic and degrees of freedom were used from the GLM for this calculation. η²p values were used to visualize the spatial distribution and relative strength of significant effects, complementing the binary significance thresholding of permutation-based p-values. False discovery rate (FDR) correction using the Benjamini-Hochberg method was applied within each set of models stratified by specific cognitive domain factor, microstructural measure, and analysis level (global whole brain summary values or regional tract-specific values). Adjusted p-values are reported for all significant effects.

When the age group × WM microstructure (global/region-based measures) interaction was significant, follow-up permutation-based GLMs were conducted within each age group separately to determine the direction and strength of associations between WM metrics and cognitive outcomes. Models included sex and years of education as covariates and used the Freedman–Lane permutation approach with 5,000 permutations to obtain p_perm_ values.

### Voxel-wise tract-based spatial statistics (TBSS) analysis

In order to assess the effect of age on white matter microstructure in a spatially unbiased manner, while complementing extracted mean summary measures and examining the effects at a finer level across the entire skeleton, whole brain voxel-wise analyses were performed using tract-based spatial statistics (TBSS; FSL v6.0.5) (Smith et al., 2006). For each diffusion modality, skeletonized maps were entered into permutation-based general linear models using FSL’s *randomise* with 5000 permutations, with age group specified as the main predictor of interest and sex included as a covariate. Statistical inference was performed using threshold-free cluster enhancement (TFCE), with significance defined at p < 0.05 (FWE-corrected). Significant clusters were identified using the thresholded statistical maps and localized by overlaying on the JHU ICBM-DTI-81 white matter labels atlas (Wakana et al., 2004). To quantify voxel-wise effects, TFCE-significant maps were binarized and summarized as (i) percent of the mean-FA skeleton covered, and (ii) median and peak |t| within the significant mask. For anatomical localization, significant masks were intersected with the JHU ICBM-DTI-81 atlas labels and labels containing ≥50 significant skeleton voxels were tabulated; unlabeled voxels (“Unclassified”) were excluded in our report.

## Results

### Demographic characteristics

An overview of the sample cohort size, demographic information, and domain specific cognitive composites at baseline visit is provided in **Table 1**. By design, age differed substantially between groups, while the distribution of sex (Females, Young: 72.7%; Older: 60.4%) and years of education (Young: 16.00 ± 3.07; Older: 17.26 ± 2.19) were broadly comparable. Both groups performed within the cognitively normal range on the Mini-Mental State Examination (MMSE, Young: 29.24 ± 1.06; Older: 28.96 ± 1.22). Baseline cognitive performance was summarized using six principal component analysis-derived composite scores representing executive function, language, working memory, verbal memory, semantic memory, and visual memory. These factor composites were approximately standardized and reported for a subset of participants with complete neuropsychological data (n = 73; Young: n = 32, Older: n = 41).

**Table 1.**
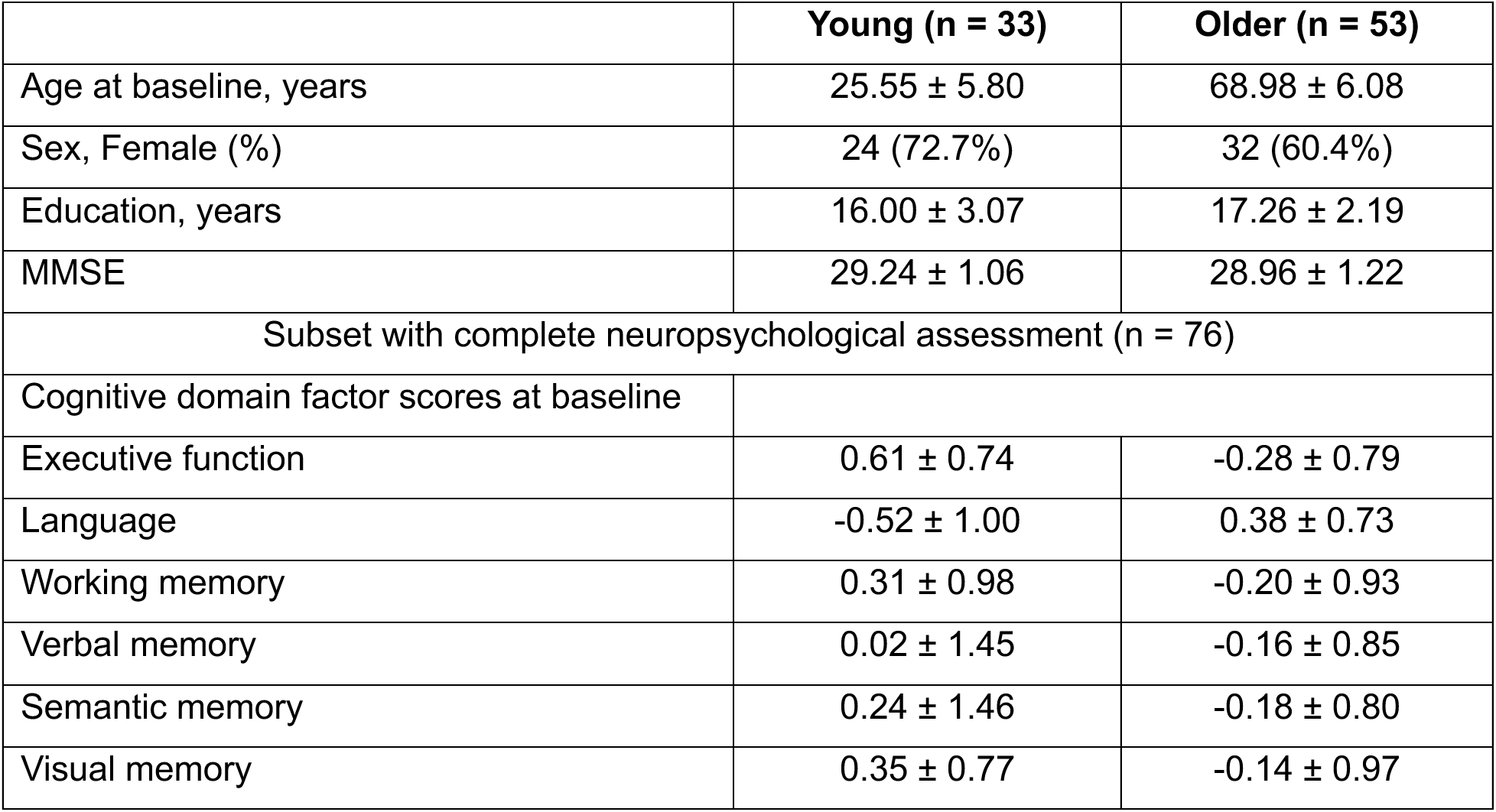
Sample cohort participant characteristics. Demographics are reported for the full cohort (n = 86). Cognitive domain factor scores are reported for the subset with complete neuropsychological data (n = 76; Young n = 32; Older n = 41). Scores were derived from PCA and are approximately standardized. Mean ± SDs are displayed for continuous variables, absolute numbers and percentages (%) are displayed for categorical variables. Abbreviations: MMSE, Mini Mental State Examination.

|  | Young (n = 33) | Older (n = 53) |
| --- | --- | --- |
| Age at baseline, years | 25.55 ± 5.80 | 68.98 ± 6.08 |
| Sex, Female (%) | 24 (72.7%) | 32 (60.4%) |
| Education, years | 16.00 ± 3.07 | 17.26 ± 2.19 |
| MMSE | 29.24 ± 1.06 | 28.96 ± 1.22 |
| Subset with complete neuropsychological assessment (n = 76) |  |  |
| Cognitive domain factor scores at baseline |  |  |
| Executive function | 0.61 ± 0.74 | -0.28 ± 0.79 |
| Language | -0.52 ± 1.00 | 0.38 ± 0.73 |
| Working memory | 0.31 ± 0.98 | -0.20 ± 0.93 |
| Verbal memory | 0.02 ± 1.45 | -0.16 ± 0.85 |
| Semantic memory | 0.24 ± 1.46 | -0.18 ± 0.80 |
| Visual memory | 0.35 ± 0.77 | -0.14 ± 0.97 |

### Associations between age and global measures of white matter microstructure

Age group effects on mean white matter skeleton measures were assessed across the five diffusion microstructural metrics within the full sample cohort (n = 86), controlling for sex. Significant main effects of age group were observed for fractional anisotropy (FA; *F* = 39.5, p_FDR_ = 3.33 × 10⁻⁴), mean diffusivity (MD; *F* = 11.4, p_FDR_ = 2.25 × 10⁻³), orientation dispersion index (ODI; *F* = 47.7, p_FDR_ = 3.33 × 10⁻⁴), and FWF (*F* = 68.4, p_FDR_ = 3.33 × 10⁻⁴), with Older adults showing lower FA and higher MD, ODI, and FWF relative to Young adults. Neurite density index (NDI) showed a trend towards lower values in Older adults (*F* = 3.46, p_FDR_ = 0.076), although this did not meet the FDR-adjusted threshold for significance. No significant main effects of sex were observed across all diffusion metrics. **Figure 1A** illustrates these group-level findings and complete statistical results are reported in **Table S1**.

**Figure 1.**
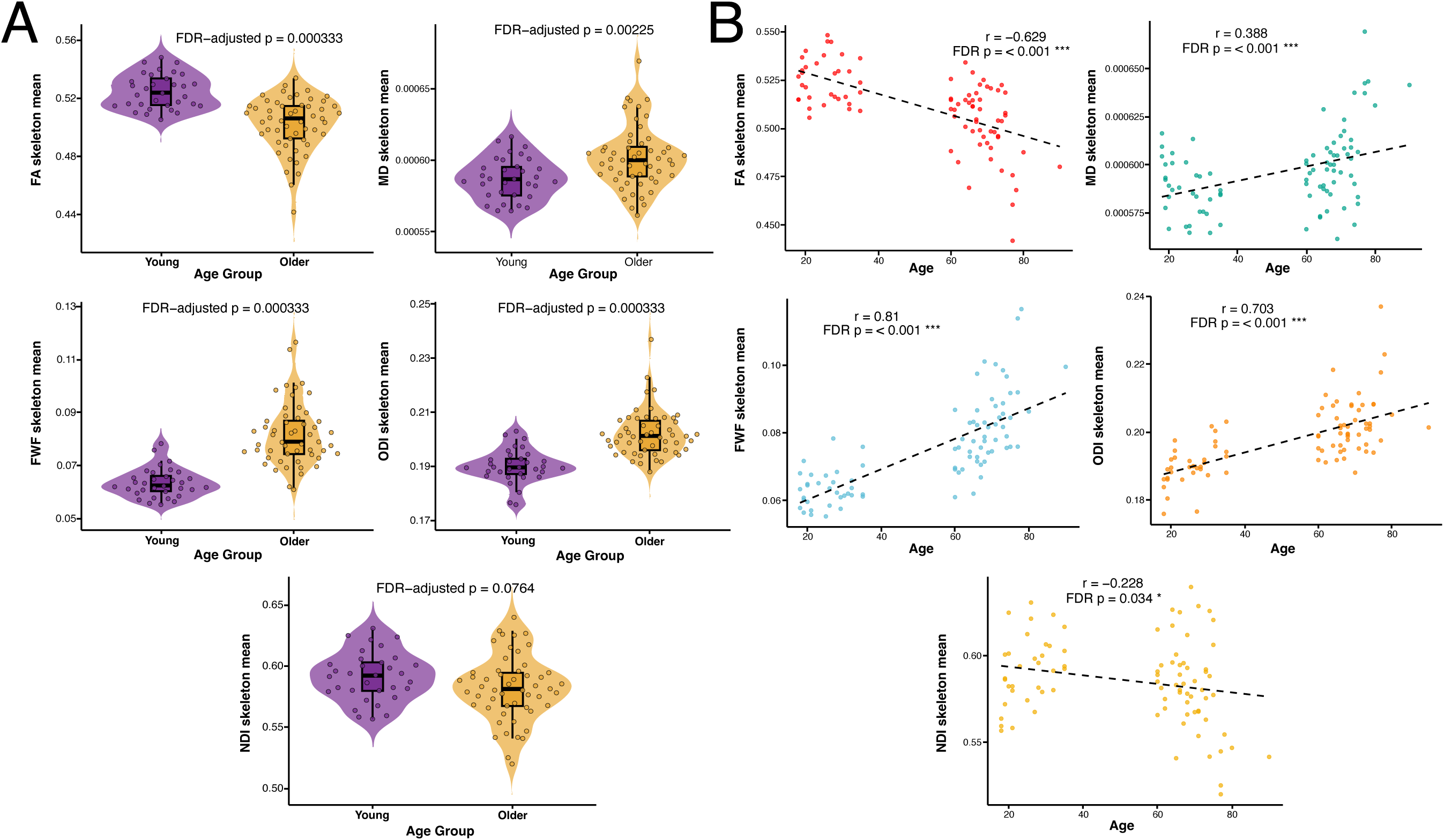
**(A**) Boxplots show age group differences (Young vs. Older) in mean WM skeleton values for five diffusion metrics, fractional anisotropy (FA), mean diffusivity (MD), neurite density index (NDI), orientation dispersion index (ODI), and free-water fraction (FWF). Group comparisons assessed with permutation-based GLMs including sex as a covariate revealed that older adults showed significantly lower FA and higher MD, ODI, and FWF, while NDI showed a negative trend in the Older group (after FDR correction). **(B**) Scatterplots display associations between chronological age (in years) and mean white matter skeleton values across the same five diffusion metrics. Strength and direction of associations are quantified with Spearman’s rank correlations (ρ, p FDR corrected). Age was associated with significantly lower FA and NDI, and higher MD, ODI, and FWF. Each shaded point represents an individual subject.

To complement these categorical age group analyses, Spearman’s rank-order correlations were computed to quantify the strength and direction of monotonic associations between chronological age (years) and mean microstructure measures across the white matter skeleton (**Figure 1B**). Older age was associated with lower FA (ρ = −0.63, p_FDR_ < 0.001) and NDI (ρ = −0.23, p_FDR_ = 0.034), and with higher MD (ρ = 0.39, p_FDR_ < 0.001), FWF (ρ = 0.81, p_FDR_ < 0.001), and ODI (ρ = 0.70, p_FDR_ < 0.001). These results overall replicated those found with age as a dichotomized variable, except that a significant association between age and white matter microstructure was also found with NDI.

### Associations between age and regional tract-based white matter microstructure

Permutation-based GLMs revealed that all 50 atlas-defined white matter tract ROIs exhibited significant age group differences (p_FDR_ < 0.05) in at least one of the five diffusion modalities (FA, MD, NDI, ODI, FWF), indicating widespread age-related alterations on brain white matter tracts. The number of tract ROIs showing significant age effects varied by modality. FWF showed the most extensive effects (48 tracts, 96%), followed by FA (42 tracts, 84%), MD (32 tracts, 64%), ODI (27 tracts, 54%), and NDI (26 tracts, 52%) (see **Figure 2B**). Effect sizes for significant age group findings were moderate to large (mean partial η² = 0.203) across tract ROIs, with the strongest overall effect observed in the fornix (FWF, ηp² = 0.509, F = 86.05, p < 0.0002). Older adults were found to have significantly lower FA values in 40 of 50 tracts (80%) and lower NDI values in 17 tracts (34%), while MD (30 tracts, 60%), ODI (27 tracts, 54%), and FWF (48 tracts, 96%) were higher in older adults.

**Figure 2.**
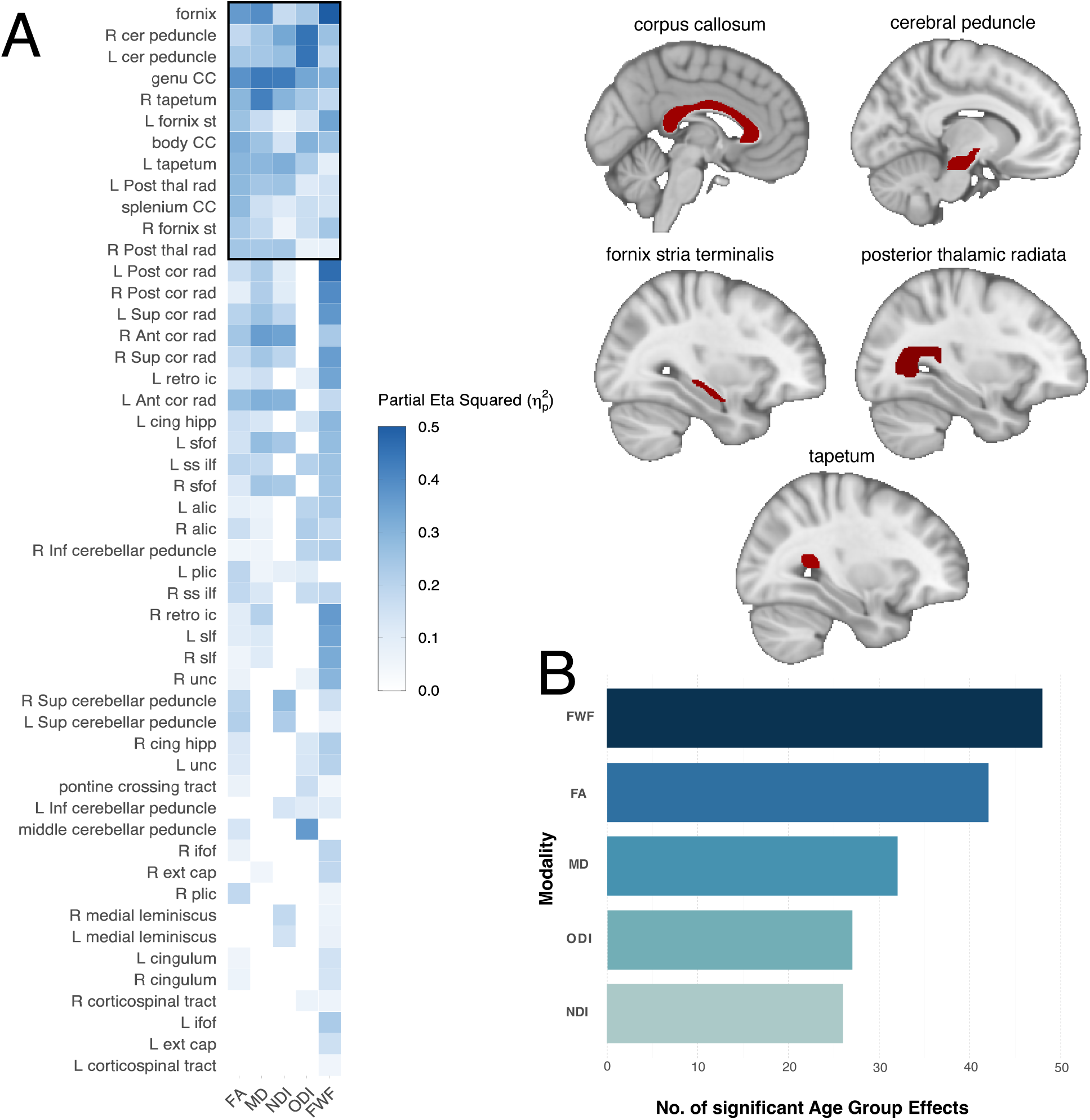
**(A)** Heatmap showing the effect size distribution of partial η² values for significant age group effects (p_FDR_ < 0.05) across 50 atlas-defined white matter tract ROIs and five diffusion modalities, FA, MD, NDI, ODI, FWF. Color intensity indicates effect size and white blank cells denote non-significance. Cells that are boxed in bold show white matter tract ROIs that are significant for age group effects across all five diffusion modalities. These 12 tracts, namely bilateral cerebral peduncle, fornix stria terminalis, posterior thalamic radiata, tapetum, as well as the fornix and corpus callosum are visualized and highlighted. **(B)** Bar graphs are grouped by modality and reflect counts of ROIs meeting the p_FDR_ < 0.05 threshold. FWF and MD tended to yield the greatest number of significant age-group differences, followed by ODI, NDI, and FA, consistent with differential sensitivity of diffusion metrics to age-related WM microstructural alterations.

Notably, a subset of twelve ROIs demonstrated significant age group effects across all five diffusion modalities (as bolded in **Figure 2A**). These respectively include the genu, body, and splenium of the corpus callosum; bilateral tapetum; bilateral fornix; bilateral stria terminalis; bilateral posterior thalamic radiation; and bilateral cerebral peduncle. Effect sizes within this set of ROIs were consistently large (ηp² range = 0.113–0.509), with the strongest observed in the fornix (FWF, ηp² = 0.509), genu of the corpus callosum (MD ηp² = 0.436; NDI ηp² = 0.433), and bilateral cerebral peduncle (ODI ηp² = 0.450–0.452). Additionally the bilateral superior fronto-occipital fasciculus (FA ηp² = 0.122–0.145; MD ηp² = 0.248–0.273; NDI ηp² = 0.236–0.241; FWF ηp² = 0.245–0.274) and sagittal stratum (FA ηp² = 0.183–0.195; MD ηp² = 0.126–0.181; ODI ηp² = 0.170–0.210; FWF ηp² = 0.186–0.255) showed significant age group effects across four modalities and the bilateral superior longitudinal fasciculus showed significant age group effects across three modalities (FA ηp² = 0.051–0.103; MD ηp² = 0.110–0.121; FWF ηp² = 0.322–0.339). Anatomically, these ROIs spanned major commissural pathways (corpus callosum, tapetum) and limbic-associated tracts (fornix, fornix stria terminalis), representing regions of multi-modality convergence where age group effects were consistently detectable across all microstructural measures. In addition, long-range association fibers, including the bilateral superior fronto-occipital fasciculus and sagittal stratum (all significant in four modalities) and the bilateral superior longitudinal fasciculus (three modalities) also showed robust multimodal effects, underscoring the involvement of association systems alongside commissural and limbic pathways. Overall, permutation-based GLMs at the individual tract ROI level revealed robust age group differences across multiple white matter tracts and across diffusion modalities (see **Figures 2A and B**).

### Voxel-wise age group differences on white matter microstructure

To capture more spatially precise age-related patterns across the white matter skeleton, TBSS analyses on the skeletonized maps were conducted for each diffusion metric. Whole brain voxel-wise analyses revealed widespread age group differences that corroborated tract ROI findings. Effect directions were consistent with ROI based results, where relative to younger adults, the older group showed significantly lower FA and NDI, and higher MD, ODI, and FWF across the white matter skeleton (**Figure S1**). In terms of spatial extent of significant age effects, FWF accounted for the largest proportion of the skeleton (47.03%), followed by FA (41.77%), MD (35.37%), NDI (23.81%), and ODI (7.91%). Interestingly, complementary voxel-wise magnitude summaries – defined as median absolute t-static within TFCE-significant voxels, revealed that ODI had the strongest local effects (median |t|=3.35), whereas effects for other diffusion-based modalities showed broader spatial coverage but lower magnitude (median |t|=2.95 for MD, median |t|=2.89 for NDI, median |t|=2.78 for FA, median |t|=2.39 for FWF). Across modalities, significant clusters were most prominent in commissural tracts such as segments of the corpus callosum, projection pathways (limbs of the internal capsule), and limbic tracts such as the fornix. Findings from the voxel-wise TBSS approach complemented results from tract ROI-level and global mean white matter measures, while providing finer-grained localization of age group related microstructural differences.

### Interaction of age group and global measures of white matter microstructure on cognition

To further assess the effect of age-related change in white matter microstructure on cognition, we performed interaction analyses that revealed statistically significant interaction effects between age group and mean white matter skeleton measures on cognitive performance. Three of the 30 tested cognitive domain (i.e., six cognitive domains) –modality (i.e., five diffusion metrics) interactions retained significance after FDR correction (**Figure 3**). These were observed for visual memory and ODI (F = 5.21, p_FDR_ = 0.022), visual memory and NDI (F = 4.70, p_FDR_ = 0.037), and for language and ODI (F = 4.21, p_FDR_ = 0.049). Post-hoc analyses stratified by age group showed that higher ODI was associated with poorer visual memory performance in the Young group (F = 7.75; p_perm_ = 0.007), but showed no significant association in the Older group (F = 0.011; p_perm_ = 0.919). Significant interaction patterns suggested that NDI was positively associated with visual memory in the Young group but not in the Older group, and that ODI was negatively associated with language in the Young group, but not in the Older group, although neither association reached significance in post-hoc age-stratified analyses.

**Figure 3.**
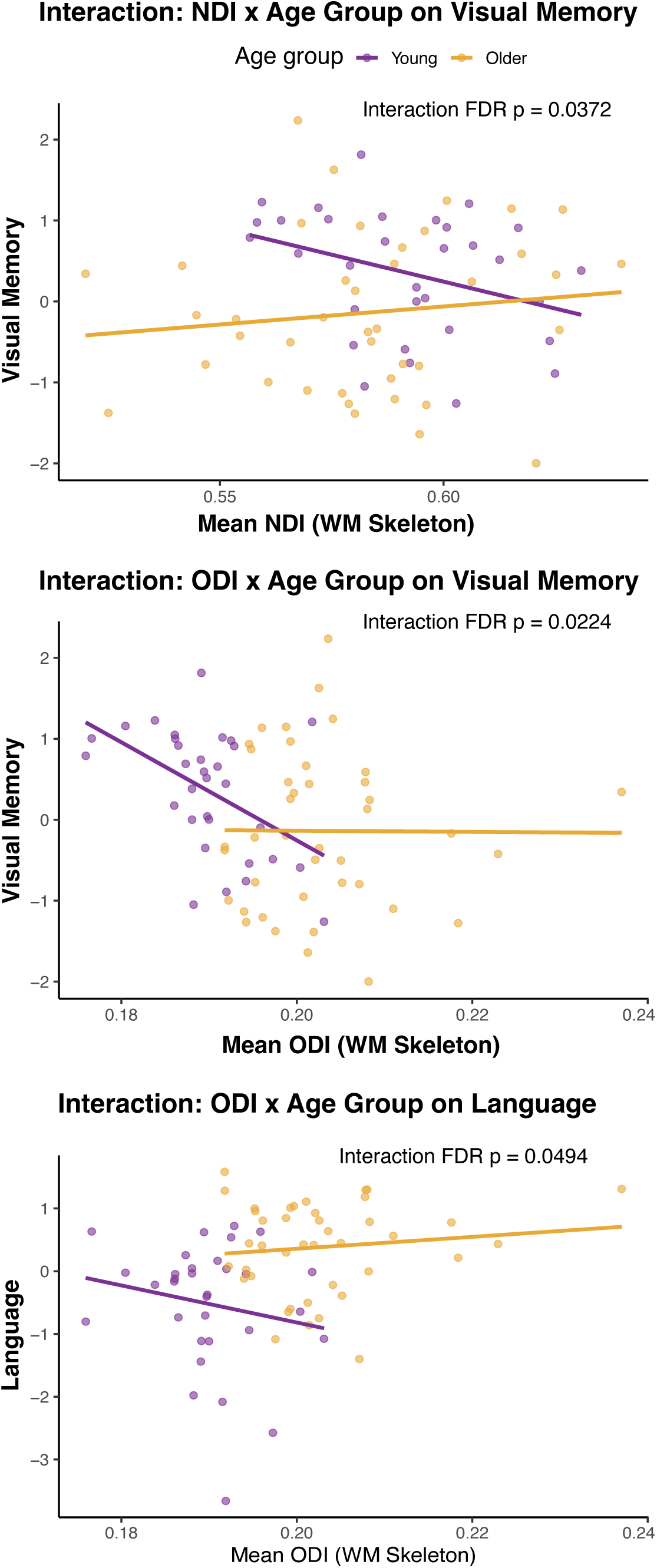
Scatterplots show significant ROI × age group interaction effects from permutation-based GLMs (controlling for sex and education, 5,000 permutations and FDR corrected). Regression lines reflect group-specific associations, with shaded points representing individual data. (i) Higher neurite density index (NDI) was associated with better visual memory in the Young group but showed little association in the Older group. (ii) Higher orientation dispersion index (ODI) was linked to significantly poorer visual memory in the Young group, with no effect in the Older group. (iii) Higher ODI predicted poorer language performance in the Young group but a weak positive association in the Older group.

### Interaction of age group and regional tract measures of white matter microstructure on cognition

Age group × regional tract microstructure interaction analyses on cognitive domains identified 80 significant interactions (p_FDR_ < 0.05) distributed across all six cognitive factor domains and five diffusion microstructural metrics. **Figure 4** illustrates an overview of these significant results. Broadly, the number of significant interactions varied by domain, with the largest counts observed for language (n = 21) and visual memory (n = 21). Across diffusion modalities, NODDI metrics made up for 65/80 (∼81%) of significant interaction effects where ODI yielded the highest number of significant interactions (n=25), followed by FWF (n=21), NDI (n=19), MD (n=9), and FA (n=6), suggesting that NODDI indices like ODI and NDI most consistently supported age-dependent coupling between regional white-matter microstructure and cognition across cognitive domains.

**Figure 4.**
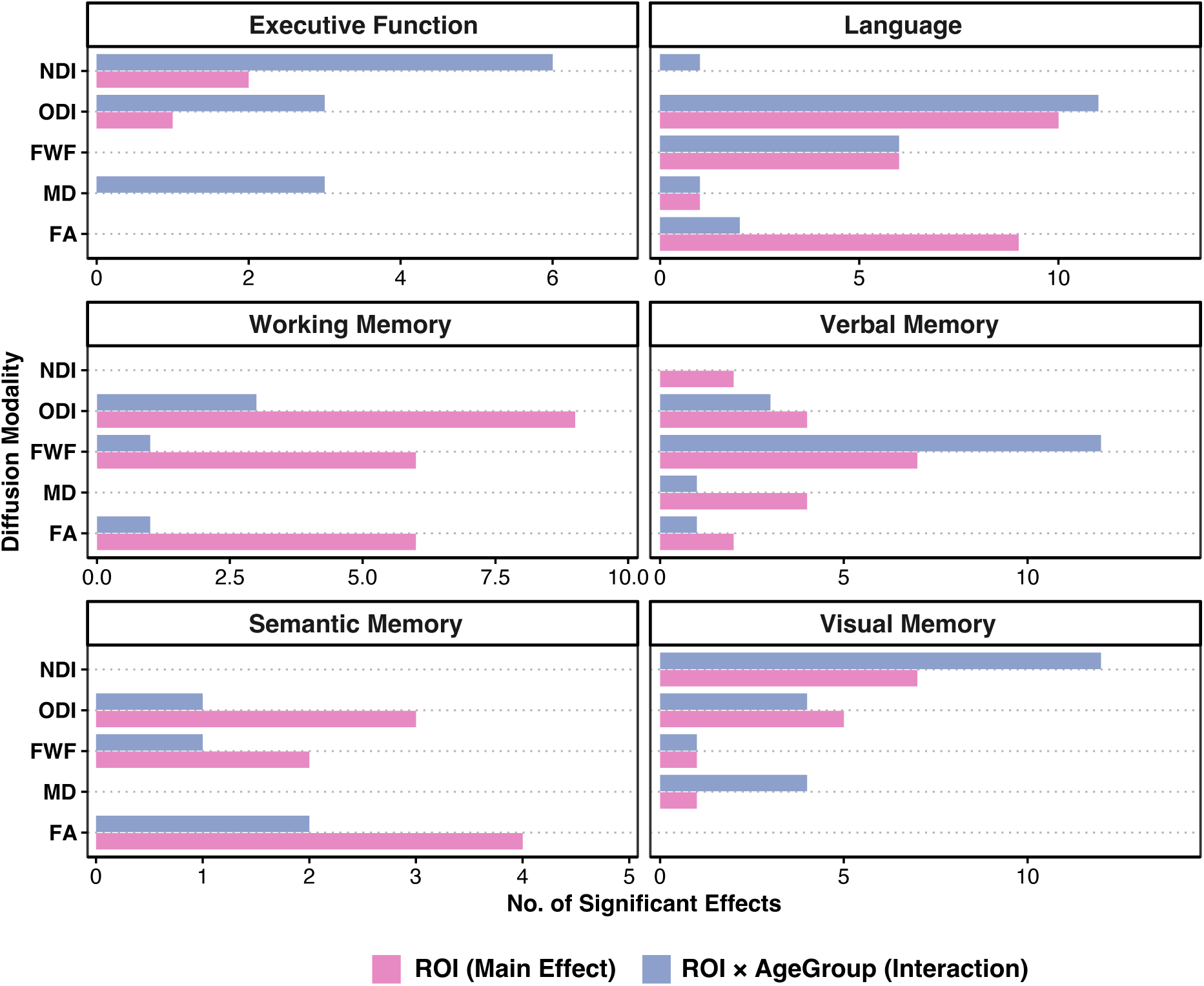
Number of significant white-matter (WM) ROI effects by diffusion modality for each cognitive domain. Bar plots display counts of atlas-defined WM tracts showing significant (p_FDR_ < 0.05) main effects of ROI (colored in pink) or ROI × age group interactions (blue) from permutation-based GLMs predicting six cognitive factor domains: executive function, language, working memory, verbal memory, semantic memory, and visual memory. Results are displayed separately for five diffusion metrics, FA, MD, ODI, NDI, FWF. Counts are aggregated across all significant effects within each modality-cognitive domain combination, providing a modality-level summary of the distribution and type of significant structure–function associations. FWF and ODI yielded the highest number of significant associations across domains, followed by NDI, MD, and FA, indicating greater sensitivity of NODDI metrics to age-related modulation of WM-cognition relationships.

Post-hoc age-stratified tests revealed that 47 of 80 significant interactions (58.8%) showed significant tract microstructure–cognition associations in at least one age group. Of these, 24 were exclusive to the Young group, 13 exclusive to the Older group, and 5 present in both groups. In the Young group, interactions were most frequent for language–ODI (n = 8), visual memory–ODI (n = 6), and visual memory–NDI (n = 4), typically involving commissural and association tracts. In the Older group, the most frequent effects occurred for executive function–NDI (n = 6) and language–NDI (n = 4), localizing to projection and limbic tracts. In the Young group, higher ODI or NDI was frequently associated with better visual memory, whereas in the Older group, higher NDI was generally associated with better executive function and language performance.

## Discussion

Aging alters white matter microstructure and the way white matter supports cognition. Here using multi-parameter diffusion MRI modeling, we show these effects are tract- and domain-specific and are most evident in NODDI-derived indices, capturing mechanistic processes beyond conventional DTI.

Extensive work in diffusion imaging have converged on a robust aging signature, demonstrating that aging is often accompanied by loss of microstructural organization, classically expressed as lower FA and higher MD (Salat et al., 2005; Madden et al., 2009), and that age-linked white matter microstructural changes are most prominent in callosal, long-range association, and limbic pathways (Head et al., 2004; Sexton et al., 2011). Yet tensor metrics conflate multiple biophysical sources and are vulnerable to extracellular partial-volume signal, motivating the use of compartment models such as NODDI to separate putative changes in neurite packing/density (NDI), fiber geometry/dispersion (ODI), and extracellular/free-water content (FWF). Consistent with prior work, we found age effects on white matter microstructural measures, where were both significant and directionally consistent, whereby older adults showed elevated MD, FWF, and ODI, accompanied by decreased FA and NDI. Within this framework, our data align with and extend these observations across analytical scales – namely, global skeleton means, atlas-based ROI tracts, and TBSS voxel-wise tests, with FWF emerging as the measure with the strongest age effects. This is supported by cohort studies suggesting FWF to be an age-sensitive biomarker which frequently shows broader or stronger age effects than FA/MD (Gullett et al., 2020; Gustavson et al., 2023; Peter et al., 2025) – in addition, free-water correction attenuates FA/MD aging differences, implying that this CSF component drives a substantial share of diffusion sensitivity to aging (Maillard et al., 2019). Voxel-wise TBSS maps mirrored these effects, showing that FWF carried the largest significant spatial proportion with diffuse age-related increases across the WM skeleton, supporting the view that extracellular compartment drives a substantial share of tensor sensitivity to underlying WM changes (Pasternak et al., 2009). In contrast, measures such as ODI showed the strongest age-related effects, but they were confined to a smaller spatial distribution, primarily within projection pathways, the callosal body and the fornix. Taken together, this dissociation between spatial extent and magnitude may offer a mechanistical account of normative white matter aging in which a diffuse extracellular process (FWF) underlies the widespread pattern observed, while packing and dispersion changes appear more localized (ODI, NDI). This may clarify why measures such as FA and MD are sensitive but mechanistically ambiguous, whereas FWF/ODI/NDI may provide greater biological specificity.

Anatomically, there was multi-modal convergence of significant age effects observed across all five WM microstructural measures in a set of bilateral tracts on an ROI level, spanning the commissural (corpus callosum segments, tapetum), limbic (fornix, stria terminalis), and projection pathways (posterior thalamic radiation [PTR], cerebral peduncle), indicating a cross-system anatomical signature associated with aging rather than a single locus. Alongside this convergent pattern, association pathways also demonstrated substantial age sensitivity; in particular, the superior longitudinal fasciculus showed significant age effects across most microstructural (4/5) measures, indicating robust vulnerability. Collectively, these findings align with existing models of WM aging in which long-range, metabolically demanding pathways with protracted developmental trajectories (Lebel et al., 2008; Grotheer et al., 2022), are disproportionately vulnerable later in life (Bartzokis, 2004). This topographic pattern of WM decline is theorized to reflect an inversion of the developmental sequence of myelination, or “last-in-first-out”, or retrogenesis, in which late-maturing tracts are more susceptible to aging processes (Stricker et al., 2009; Brickman et al., 2012). Population-scale cellular mapping of late-life WM microstructure further supports this similar view whereby association, commissural, and limbic tracts that show steep age-related decline also exhibit enrichment for oligodendrocyte and glial gene expression programs, suggesting that their vulnerability reflects shifts in myelin maintenance and glial support beyond purely axonal loss (Chin et al., 2025). In the context of the present findings, this convergence suggests that tract types showing multimodal age sensitivity may be those in which age-related microstructural changes are most strongly coupled to alterations in myelin maintenance and glial support, rather than reflecting axonal loss alone. Consistent with this broader organizational gradient, we also observed tract-specific signatures consistent with prior work. In the corpus callosum which is heterogeneous in its microstructural composition (Aboitiz et al., 1992), age effects were also markedly larger in the genu/body than splenium across diffusion modalities, consistent with the hypothesis of an anterior-to-posterior gradient in callosal aging (Ota et al., 2006; Kraft et al., 2024). The fornix, which showed the strongest age effect, has been established to be a vulnerable limbic pathway and early marker for medial temporal lobe atrophy (Oishi et al., 2012). Furthermore, large-scale aging cohort studies (e.g. UK Biobank) have indicated that thalamic radiations such as the PTR also show pronounced age-related effects (Cox et al., 2016). Therefore, our findings provide evidence supporting age-associated white matter changes, lending novel findings showing that aging-related changes in white matter are associated with variation across multiple microstructural properties, including those linked to glial-related tissue features, in a regionally selective manner.

Having established the microstructural profile of aging, we then assessed whether age also influences the relationship between brain structure and cognition, i.e. whether microstructure properties of white matter tracts also show age-dependent links to cognitive domains. We found that age moderates WM-cognition relationship, and that the pattern was metric- and tract type-specific. At the global WM skeleton level, higher NDI was significantly associated with lower visual memory and higher ODI was significantly associated with lower language and visual memory in younger adults; these associations were attenuated in older adults. ODI has been previously linked with aspects of cognition in older adults as an index of orientation dispersion/loss of fiber coherence (Raghavan et al., 2021), higher values in younger adults could imply microstructural disorganization and less efficiency in information integration underlying cognitive domains (Koirala et al., 2021). Although higher NDI is expected to better support cognition, inverse patterns have been reported in younger/healthy cohorts, whereby better cognitive performance has been linked to more efficiently pruned pathways (reflected as lower NDI) that refines long-range signaling rather than a single narrative that “more tissue/packing” is uniformly better. When observed in tandem, it has also been suggested the pattern of high coherence (ODI) and lower density (NDI) within a younger cohort could imply smaller and less dispersed axonal organization reflecting microstructural architecture that restricts synaptic connections to a cost-effective minimum and facilitates neural efficiency (Koirala et al., 2021). The age-related slope attenuation for NDI and ODI metrics in the Older group may be interpreted through the lens of neural dedifferentiation and compensation/scaffolding frameworks which posit that aging is accompanied by less tightly coupled, specific structure-behavior mappings, while shifting performance support to alternate, more distributed circuitries and neuroplasticity (Lövdén et al., 2010; Reuter-Lorenz and Park, 2014; Koen and Rugg, 2019).

It is intriguing that, on a tract-based ROI scale, age dependent WM-cognition interactions were driven predominantly by NODDI indices, with ODI and NDI accounting for the larger share of these effects, indicating coherence/dispersion and neurite packing are the principal axes along which aging alters structure-behavior relationships. Consistent with global WM skeleton findings, younger adults showed stronger NDI and ODI associations with language and visual memory, especially within association and callosal pathways. With aging, the structure-behavior topography shifts towards limbic and projection conduits, where older adults showed stronger links between NDI and pathways such as the fornix and internal capsule for executive function and language. Notably, although FWF demonstrated high significant interaction effects, these effects showed limited significant WM-cognition associations within each age group. This apparent dissociation may reflect the dual role of free water as a general marker of tissue milieu changes, highly sensitive to between-group aging differences, yet less proximally tied to tract-specific cognitive variance within stratified age groups. Biophysically, FWF captures the proportion of isotropic extracellular water arising from neuroinflammatory, microvascular and atrophic processes which are known to increase with age and are associated with general age-group differences in cognition at a systems level (Pasternak et al., 2009). In this sense, FWF complements the other NODDI indices such as ODI and NDI, which capture coherence/dispersion and neurite packing/volume, by providing a broader measure of WM vulnerability and tissue health, thereby contextualizing microstructural changes more intrinsically tied to tract-specific cognitive operations.

While the present framework captures WM variability in a healthy aging sample, our NODDI-based findings on white matter aging and its age group-specific impact on cognition have significant implications for neurodegenerative or neurological disorders, such as Alzheimer’s disease and mild traumatic brain injury (mTBI), which affect white matter microstructure and cognitive function. Among patients with mTBI, a known risk factor of developing Alzheimer’s disease, Palacios et al. (2020) found that, cross-sectionally, early decreases of FA and increases of MD in correspondence to elevated FWF in extensive WM tracts were reported in mTBI patients, while longitudinal changes in WM were more pronounced for NDI. These results suggest that NODDI is a more sensitive and specific biomarker than DTI for WM microstructural changes in mTBI and may advance diagnosis, prognosis, and treatment monitoring in this clinical population. In addition, recent studies found differential longitudinal trajectories in WM changes between normal and abnormal aging across the continuum of Alzheimer’s disease, demonstrating tract-specific accelerated microstructural aging measured by NODDI (Archer et al., 2023). Given that findings from studies in clinical populations may reflect disease-specific processes, our results provide a baseline characterization of age-related susceptibility to white matter changes across the entire white matter network. This foundation help contextualize and interpret additional disease-specific alterations and their impact on cognition.

The present analyses advance a multi-parameter, *in vivo* account of white matter aging and its age-dependent coupling to aspects of cognition. However, these findings should be interpreted in light of several limitations. First, the cross-sectional design and modest sample size limit causal inference and precision of interaction estimates. Future work is warranted by extending these analyses longitudinally to track individual trajectories and further evaluate temporal coupling between changes in structural white matter and cognition. In addition, *in vivo* microstructural indices remain as model-based approximations that conflate multiple biophysical processes rather than singularly reflecting any one cellular mechanism or pathology, thus interpretation remain indirect and should be triangulated with histological or multimodal data (for example, PET or myelin-sensitive MRI). Despite these limitations, the present study using MRI-based NODDI modeling provides evidence for biophysically distinct microstructural processes underlying age-related white matter and cognitive changes even with a relatively modest sample size, in a non-invasive manner, while providing accessible and cost-effective research and clinical utility for aging and age-related neurodegenerative disorders.

The current work highlights the microstructural heterogeneity of white-matter aging, revealing that incorporating alternate diffusion models such as NODDI offer enhanced sensitivity to age-related alterations and their cognitive correlates. Across analytic scales, we observed age-linked, diffuse increases in free water and more spatially focused changes in fiber dispersion and density, with cognitive associations strongest for NODDI measures. Collectively, these findings establish a neurobiologically grounded framework for understanding the relationship between microstructural alterations in white matter and age-related differences in cognition.

## Author Contributions

RC and HO designed and supervised the research. RC, XH, JR, J-HK, HO performed the research. RC, XH, JR, J-HK, and HO provided analytic support and interpretation of the results. RC and HO wrote the manuscript and made figures. All authors revised the manuscript.

---

**Table S1.**
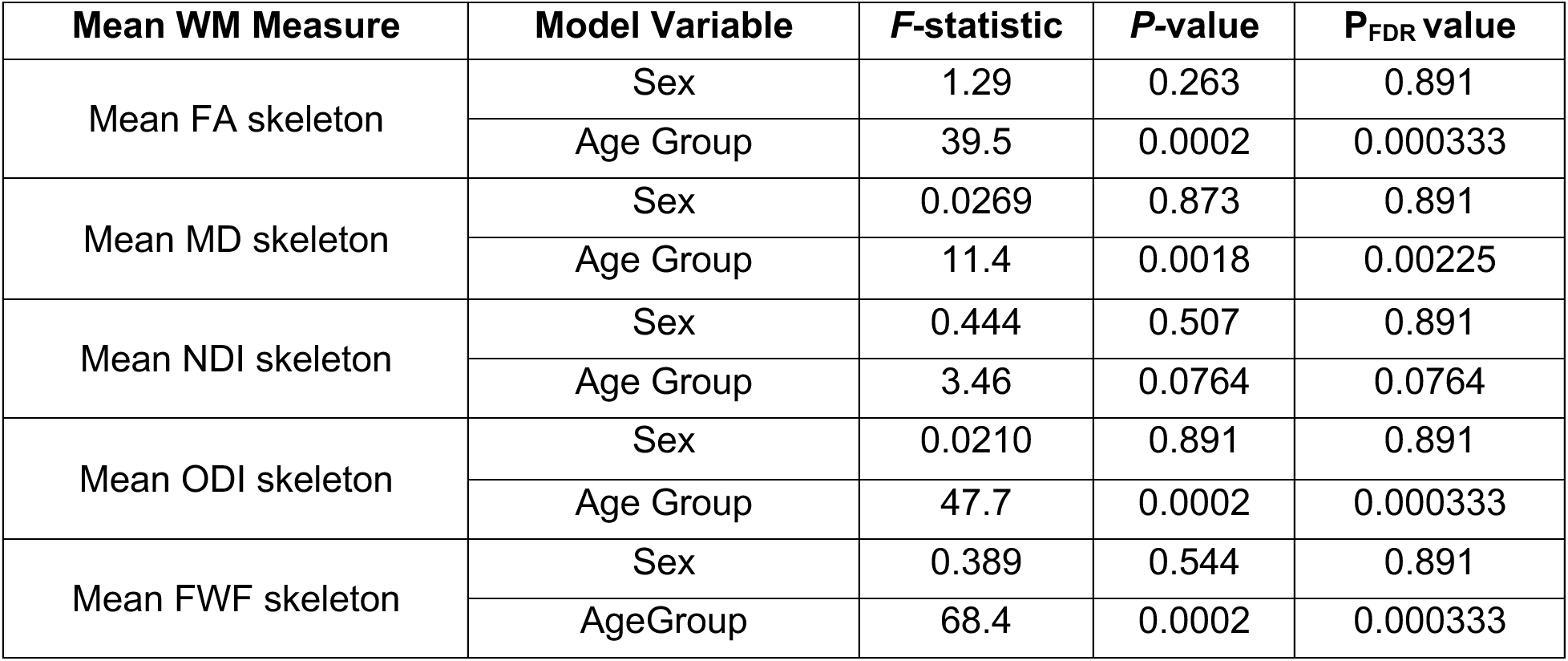
Permutation-based general linear model results for main effects of age group and sex on mean white matter skeleton microstructure measures (N = 86).

| Mean WM Measure | Model Variable | *F*-statistic | *P*-value | *P*<sub>FDR</sub> value |
| --- | --- | --- | --- | --- |
| Mean FA skeleton | Sex | 1.29 | 0.263 | 0.891 |
|  | Age Group | 39.5 | 0.0002 | 0.000333 |
| Mean MD skeleton | Sex | 0.0269 | 0.873 | 0.891 |
|  | Age Group | 11.4 | 0.0018 | 0.00225 |
| Mean NDI skeleton | Sex | 0.444 | 0.507 | 0.891 |
|  | Age Group | 3.46 | 0.0764 | 0.0764 |
| Mean ODI skeleton | Sex | 0.0210 | 0.891 | 0.891 |
|  | Age Group | 47.7 | 0.0002 | 0.000333 |
| Mean FWF skeleton | Sex | 0.389 | 0.544 | 0.891 |
|  | AgeGroup | 68.4 | 0.0002 | 0.000333 |

**Figure S1.**
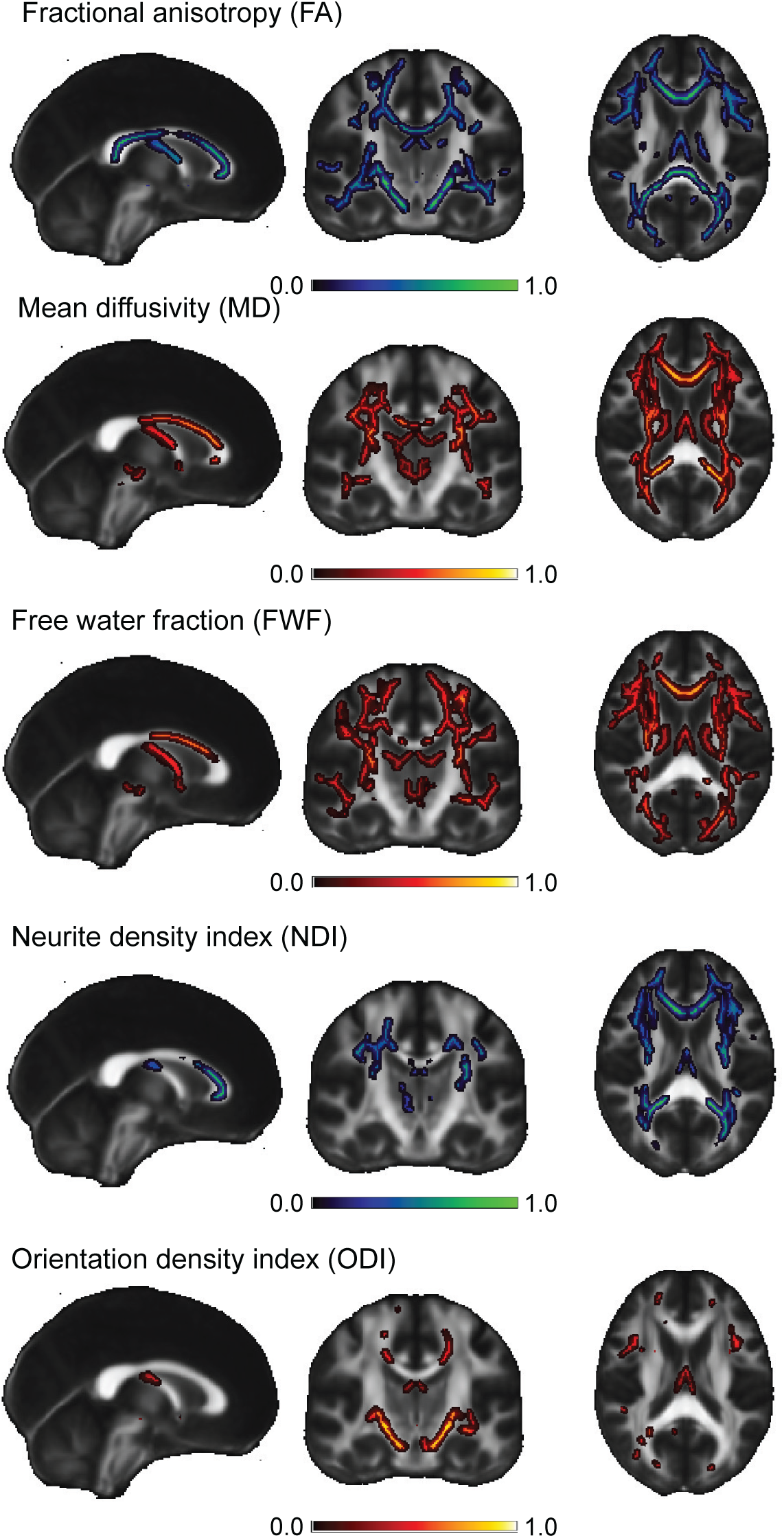
Voxel-wise analyses using tract-based spatial statistics (TBSS) revealed widespread age group related differences in WM microstructure across diffusion metrics. Results are thresholded at TFCE-corrected p < 0.05 and displayed on the MNI152 T1 template (1mm) with the group-mean FA skeleton. FA and NDI show regions where Older < Younger effects, whereas MD, FWF, and ODI show Older > Younger effects.

